# *PKD2* influence uric acid levels and gout risk by interacting with *ABCG2*

**DOI:** 10.1101/278713

**Authors:** Zheng Dong, Jingru Zhou, Shuai Jiang, Yuan Li, Dongbao Zhao, Chengde Yang, Yanyun Ma, Hongjun He, Hengdong Ji, Li Jin, Hejian Zou, Jiucun Wang

## Abstract

**Background:** Uric acid is the final product of purine metabolism and elevated serum urate levels can cause gout. Conflicting results were reported for the effect of *PKD2* on serum urate levels and gout risk. Therefore, our study attempted to state the important role of *PKD2* in influencing the pathogenesis of gout.

**Method:** SNPs in *PKD2* (rs2725215 and rs2728121) and *ABCG2* (rs2231137 and rs1481012) were tested in approximately 5,000 Chinese individuals.

**Results:** Two epistatic interactions between loci in *PKD2* (rs2728121) and *ABCG2* (rs1481012 and rs2231137) showed distinct contributions to uric acid levels with *P* _int_ values of 0.018 and 0.004, respectively, and the associations varies by gender and BMI. The SNP pair of rs2728121 and rs1481012 justly played roles in uric acid in females (*P* _int_ = 0.006), while the other pair did in males (*P* _int_ = 0.017). Regarding BMI, the former SNP pair merely contributed in overweigh subjects (*P* _int_ = 0.022) and the latter one did in both normal and overweigh individuals (*P* _int_ = 0.013 and 0.047, respectively). Furthermore, the latter SNP pair was also associated with gout pathology (*P* _int_ = 0.001), especially in males (*P* _int_ = 0.001). Finally, functional analysis showed potential epistatic interactions in those genes region and *PKD2* mRNA expression had a positive correlation with *ABCG2*’s (r = 0.743, *P* = 5.83e-06).

**Conclusion:** Our study for the first time identified that epistatic interactions between *PKD2* and *ABCG2* influenced serum urate concentrations and gout risk, and *PKD2* might affect the pathogenesis from elevated serum urate to hyperuricemia to gout by modifying *ABCG2*.

## Introduction

Uric acid is the end breakdown product of purine complex in human beings and high uric acid levels in the blood (hyperuricemia) leads to the deposition of monosodium urate (MSU) crystals in the joints and tissues, which plays a predominant role in the development of gout (Dehghan et al. 2008; Yang et al. 2010; Köttgen et al. 2012; Perez-Ruiz et al. 2014; Dong et al. 2017b). Gout is one acute inflammatory arthritis (Dong and Wang 2015; Dong et al. 2017a) and affects a large number of individuals in many countries (Zhu et al. 2011; Dong et al. 2015a). Hyperuricemia is able to cause gout (Dong et al. 2015b), but not all hyperuricemia patients develop gout (Merriman 2015). Generally, only a quarter of hyperemia patients finally develop gout, indicating that hyperuricemia is necessary but not sufficient for gout (Merriman 2015). Nowadays, several loci in different genes have been identified to affect serum urate concentration and gout risk (Köttgen et al. 2012; Merriman 2015). However, those loci only explained a fraction of heredity of serum urate/gout and some of them contribute differently to the pathogenesis from elevated serum urate to hyperuricemia to gout (Köttgen et al. 2012; Dong et al. 2017b). And epistasis had been proved to affect complex traits and played important roles in various biological functions (Cordell 2002; Phillips 2008). Therefore, it is urgent to test the effect of epistatic interactions of genes on the development from elevated serum urate to hyperuricemia to gout.

In addition, recent studies found *PKD2* associated with uric acid and gout (Zhang et al. 2016), while other researchers had an opposite opinion (Dehghan et al. 2008). The conflicting results for the effect of *PKD2* on serum urate levels and gout risk is a challenge topic which need more evidences to obtain a reliable conclusion. In addition, a suggestive epistatic interaction for regulating serum urate concentrations had been reported in the gene region of *PKD2* and *ABCG2* (Wei et al. 2014). Therefore, epistasis analysis of *PKD2* and *ABCG2* might obtain a new evidence for this conflict and provide an explanation to extra heredity of serum urate and gout.

In the present study, we aim to study epistasis between *PKD2* and *ABCG2* applying a four-step approach. Firstly, the effect of loci interactions on serum urate was studied. Then the candidate urate-related interactions were tested in hyperuricemia and gout patients. Other heterogeneity factors, including gender, body mass index (BMI) and smoking status, influence on above significant associations were also considered in our analysis. In addition, further functional analysis was performed to provide more proof about the interactions. Finally, the relation of mRNA expression between *PKD2* and *ABCG2* were investigated. Through this strategy, the present study for the first time found that epistatic interactions between *PKD2* and *ABCG2* contributed to serum urate concentrations and gout risk, and *PKD2* might influence the pathogenesis from elevated serum urate to hyperuricemia to gout by modifying *ABCG2*.

## Methods and materials

### Study subjects

The present study was approved by the Ethical Committees of the School of Life Sciences of Fudan University (approval number of 140) and was conducted in accordance with the principles of the Declaration of Helsinki. All subjects in this study provided their written informed. All of 582 gout patients were diagnosed with gout (OMIM: #138900) following the American College of Rheumatology diagnostic criteria (Wallace et al. 1977), came from Changhai Hospital, Taixing People’s Hospital and Taizhou People’s Hospital. And gout patients did not use any urate-lowering drugs two weeks before sample collection.

Furthermore, 4332 individuals with no history of gout were selected from the Taizhou Longitudinal Study (Wang et al. 2009). Those subjects were divided into subgroups based on their smoking status recorded in questionnaires, and body mass index (BMI) values according to the categories of the World Health Organization (WHO) (Yang et al. 2016; Dong et al. 2017b). Smoking status contained non-smoker, former smoker and current smoker. Regarding BMI, three subgroups (underweight: BMI < 18.5; normal weight: 18.50 ≤ BMI < 25; overweight: BMI ≥ 25) were divided in this study. Among them, 1387 subjects were treated as hyperuricemia patients due to their high serum urate levels (> 417 umol/L) (Khanna et al. 2012), and the others were treated as healthy controls. The detail characteristics of the participants in this study are illustrated in Supplementary Material, Table S1.

### SNP genotype and Real-time qPCR

Peripheral blood was collected from all participants in this study and DNA was extracted from blood samples. RNA was isolated from blood cells of 58 male healthy subjects. Detail information of experimental approaches for DNA and RNA had been described in our previous study (Dong et al. 2017b). Genotyping of SNPs in *PKD2* (rs2725215 and rs2728121) and *ABCG2* (rs2231137 and rs1481012) was processed by SNaPshot. And the gene relative expression was tested using real-time qPCR.

### Bioinformatics analysis

Two *PKD2* SNPs sets were used in this study. One (set one) was SNPs associated with serum urate reaching genome-wide significance levels in a recent genome-wide association study (Köttgen et al. 2012). Another (set two) was all SNPs in the gene region of *PKD2*. Cell-type-specific enhancer enrichment analysis was performed in both two sets using HaploReg (http://archive.broadinstitute.org/mammals/haploreg/haploreg.php) (Ward and Kellis 2012). SNPnexus (Dayem Ullah et al. 2013) and UCSC genome browser (Kent et al. 2002) were applied for the functional annotation of loci. Enlight (http://enlight.wglab.org) was used to draw regional plot and show epigenetic modification (Guo et al. 2015). In addition, linkage disequilibrium (LD) relationships among target SNPs were tested and generated LD plots using SNAP (http://archive.broadinstitute.org/mpg/snap/index.php) (Johnson et al. 2008).

### Statistical analysis

*P*-value of SNP pair affected on serum urate concentrations was calculated by linear regression adjusted age, gender and target SNPs. Variance explained of SNP pair in serum urate was calculated by linear regression. And *P*-value of SNP pair in hyperuricemia and gout were calculated by logistic regression adjusted age, gender and target SNPs. Furthermore, subgroups analysis of gender, BMI and smoking status were also performed. In addition, the mRNA expression correlation between *PKD2* and *ABCG2* was analyzed in our study. *P* values less than 0.05 were considered statistically significant and all statistical analyses were processed by using R (Version 3.0.2: www.r-project.org/).

## Results

### Epistatic interactions of ***PDK2* and *ABCG2* contributed to serum urate concentrations**

A total of four SNP pairs were tested in our study. Of them, two SNP pairs (rs2728121:rs1481012 and rs2728121:rs2231137) were identified to associate with the concentrations of serum urate (Estimate = -14.487, *P* _int_ = 0.018 and Estimate = 9.781, *P* _int_ = 0.004, respectively), while other pairs did not (*P* _int_ = 0.181 and *P* _int_ = 0.270, respectively) (Table 1). The former SNP pair (rs2728121:rs1481012) could explain 0.099% of uric acid variance without conditioning on the marginal SNPs, and the later pair (rs2728121:rs2231137) could explain 0.164% of the variance (Table 1).

**Table 1:** Association between serum urate and epistatic interactions of loci in ***PKD2* and *ABCG2***

| Chr. | SNP<sub>1</sub> | Pos<sub>1</sub> | A<sub>1</sub> | SNP<sub>2</sub> | Pos<sub>2</sub> | A<sub>2</sub> | Dist | LD ($r^2$) | | Estimate | $P_{\text{int}}$ | Variance explained (%) |
| --- | --- | --- | --- | --- | --- | --- | --- | --- | --- | --- | --- | --- |
| 4 | rs2725215 | 88961571 | T | rs1481012 | 89039082 | G | 77511 | 0.605 | Male | -2.309 | 0.693 | - |
|  |  |  |  |  |  |  |  |  | Female | -12.817 | 0.236 | - |
|  |  |  |  |  |  |  |  |  | Total | -7.020 | 0.181 | - |
| 4 | rs2725215 | 88961571 | T | rs2231137 | 89061114 | T | 99543 | 0.116 | Male | 4.830 | 0.618 | - |
|  |  |  |  |  |  |  |  |  | Female | 20.178 | 0.220 | - |
|  |  |  |  |  |  |  |  |  | Total | 9.306 | 0.270 | - |
| 4 | rs2728121 | 88997102 | C | rs1481012 | 89039082 | G | 41980 | 0.370 | Male | -6.051 | 0.379 | - |
|  |  |  |  |  |  |  |  |  | Female | -33.315 | 0.006 | 0.493 |
|  |  |  |  |  |  |  |  |  | Total | -14.487 | 0.018 | 0.099 |
| 4 | rs2728121 | 88997102 | C | rs2231137 | 89061114 | T | 64012 | 0.045 | Male | 8.980 | 0.017 | 0.144 |
|  |  |  |  |  |  |  |  |  | Female | 9.250 | 0.171 | - |
|  |  |  |  |  |  |  |  |  | Total | 9.781 | 0.004 | 0.164 |

Chr, chromosome of a SNP pair. SNP<sub>1</sub> (SNP<sub>2</sub>), Pos<sub>1</sub> (Pos<sub>2</sub>), name and position of the first (second) SNP. A<sub>1</sub>, effect allele of SNP<sub>1</sub>; A<sub>2</sub>, effect allele of SNP<sub>2</sub>. dist, distance in bp between two SNPs. LD ( $r^2$ ), linkage disequilibrium between two SNPs. $P_{\text{int}}$ , $P$ -value of the interaction between SNP pair in serum urate was calculated by linear regression adjusted age, gender and target SNPs. Variance explained of the interaction between SNP pair in serum urate was calculated by linear regression.

In addition, the present study showed that the significant association between SNP pairs and serum urate were modified by gender (Table 1). In males, the latter interaction pair exhibited an effect on uric acid (Estimate = 8.980, *P* _int_ = 0.017) while the former interaction pair did not (*P* _int_ = 0.379). In contrast to female subjects, the former interaction pair contributed to the concentrations of serum urate (Estimate = -33.315, *P* _int_ = 0.006) while the latter interaction pair did not (*P* _int_ = 0.171). Interestingly, the explanation of former interaction pair on uric acid variance achieved 0.493% in female, suggesting an important influence of epistasis between *PKD2* and *ABCG2* on serum urate.

### Epistatic interactions of *PDK2* and *ABCG2* affected gout risk

Because elevated serum urate is a critical risk factor for the development of hyperuricemia and gout. We further checked the contributions of epistatic interactions of *PDK2* and *ABCG2* on hyperuricemia and gout. Regards hyperuricemia, the former interaction pair justly affected the hyperuricemia risk in females (Estimate = -0.648, *P* _int_ = 0.017) (Table 2), but did not in males (*P* _int_= 0.483). In opposite of the former pair, the latter pair associated with hyperuricemia in males with *P* _int_ value of 0.018, but did not in females (*P* _int_ = 0.427). With combination of males and females, the latter pair also influenced the pathogenesis of hyperuricemia (Estimate = 0.193, *P* _int_ = 0.009). For the development from hyperuricemia to gout, the latter pair played an significant role in this translation (Estimate = 0.313, *P* _int_ = 0.035), especially in males (Estimate = 0.338, *P* _int_ = 0.030) (Table 2). However, the former pair showed not significant association with this translation in both subgroup of male and female (*P* _int_ = 0.444 and *P* _int_ = 0.577, respectively). Due to the contribution of the latter pair in the pathogenesis from elevated serum urate to hyperuricemia to gout, it distinctly affected the risk of gout (Estimate = 0.480, *P* _int_ = 0.001), especially for male individuals (Estimate = 0.524, *P* _int_ = 0.001).

**Table 2:** Association between SNP pairs and hyperuricemia/gout

| SNP<sub>1</sub> | SNP<sub>2</sub> |  | <u>Hyperuricemia Vs. Control</u> |  |  | <u>Gout Vs. Hyperuricemia</u> |  |  | <u>Gout Vs. Control</u> |  |  |
| --- | --- | --- | --- | --- | --- | --- | --- | --- | --- | --- | --- |
|  |  |  | Estimate | SE | *P*<sub>int</sub> | Estimate | SE | *P*<sub>int</sub> | Estimate | SE | *P*<sub>int</sub> |
| rs2728121 | rs1481012 | Male | -0.108 | 0.153 | 0.483 | 0.188 | 0.245 | 0.444 | 0.077 | 0.258 | 0.765 |
|  |  | Female | -0.648 | 0.272 | 0.017 | -0.528 | 0.947 | 0.577 | -0.897 | 0.835 | 0.282 |
|  |  | Total | -0.254 | 0.130 | 0.051 | 0.165 | 0.234 | 0.482 | -0.004 | 0.239 | 0.987 |
| rs2728121 | rs2231137 | Male | 0.206 | 0.087 | 0.018 | 0.338 | 0.156 | 0.030 | 0.524 | 0.161 | 0.001 |
|  |  | Female | 0.118 | 0.148 | 0.427 | 0.100 | 0.536 | 0.852 | 0.229 | 0.523 | 0.661 |
|  |  | Total | 0.193 | 0.074 | 0.009 | 0.313 | 0.149 | 0.035 | 0.480 | 0.149 | 0.001 |

SNP<sub>1</sub> (SNP<sub>2</sub>), name of the first (second) SNP. SE, standard error. *P*<sub>int</sub>, *P*-value of the interaction between SNP pair in hyperuricemia and gout were calculated by logistic regression adjusted age, gender and target SNPs.

### Association between epistatic interactions and serum urate in BMI and smoking subgroups

Our previous studies have suggested that body mass index (BMI) and cigarette smoking could affect uric acid levels (Dong et al. 2015b, 2017b; Yang et al. 2016), while their contributions on the association between epistasis and serum urate was elusive. Therefore, we performed further analysis in the BMI and smoking status subgroups.

After analyzed in BMI subgroups, the latter pair was identified to be associated with uric acid in normal subjects (Estimate = 11.456, *P* _int_ = 0.013) and overweight individuals (Estimate = 9.844, *P* _int_ = 0.047) (Table 3). And the former pair only affected the concentrations of serum urate in overweight subgroup (Estimate = -21.702, *P* _int_ = 0.022). However, no significant associations were found in the subgroup of underweight with all *P*-values larger than 0.05. Regards smoking status, both SNP pairs did not influence uric acid levels (Table 3).

**Table 3:** Association between SNP pairs and serum urate in subgroups of BMI and smoking status

| SNP<sub>1</sub> | SNP<sub>2</sub> |  | <u>Subgroup-1</u> |  |  | <u>Subgroup-2</u> |  |  | <u>Subgroup-3</u> |  |  |
| --- | --- | --- | --- | --- | --- | --- | --- | --- | --- | --- | --- |
|  |  |  | Estimate | SE | *P*<sub>int</sub> | Estimate | SE | *P*<sub>int</sub> | Estimate | SE | *P*<sub>int</sub> |
| rs2728121 | rs1481012 | BMI | 19.794 | 41.811 | 0.637 | -10.483 | 8.306 | 0.207 | -21.702 | 9.477 | 0.022 |
|  |  | Smoke | -25.467 | 15.210 | 0.095 | -7.751 | 10.478 | 0.460 | -9.196 | 8.343 | 0.271 |
| rs2728121 | rs2231137 | BMI | 24.512 | 22.948 | 0.288 | 11.456 | 4.620 | 0.013 | 9.844 | 4.962 | 0.047 |
|  |  | Smoke | 9.972 | 8.385 | 0.235 | 7.106 | 5.223 | 0.174 | 7.592 | 4.608 | 0.100 |

SNP<sub>1</sub> (SNP<sub>2</sub>), name of the first (second) SNP. SE, standard error. *P*<sub>int</sub>, *P*-value of the interaction between SNP pair in serum urate was calculated by linear regression adjusted age, gender and target SNPs. Subgroup of BMI: 1, Underweight (BMI < 18.5); 2, Normal (18.5 ≤ BMI < 25); 3, Overweight (BMI ≥ 25). Subgroup of smoke: 1, non-smokers; 2, former smokers; 3, current smokers.

### Functional analysis for identified SNP pairs

Chromatin states analysis was performed in the gene region of *PKD2* and *ABCG2*, showing several strong and weak enhancers located at this region, especially at the gene region of *PKD2* (Supplementary material Figure 1 and 2). Consistent with this result, many other functional factors, such as transcription factor and DNase factor, were also identified in *PKD2* gene region (Supplementary material Figure 1 and 2). Our results suggested a potential regulate effect in those genes region, especially in *PKD2*. In addition, *ABCG2* had been identified with function as urate transporter in previous studies (Woodward et al. 2009; Köttgen et al. 2012; Dong et al. 2015a). Above all, we hypothesized that *PKD2* influenced serum urate by epistatic interacting with *ABCG2*.

**Figure 1.**
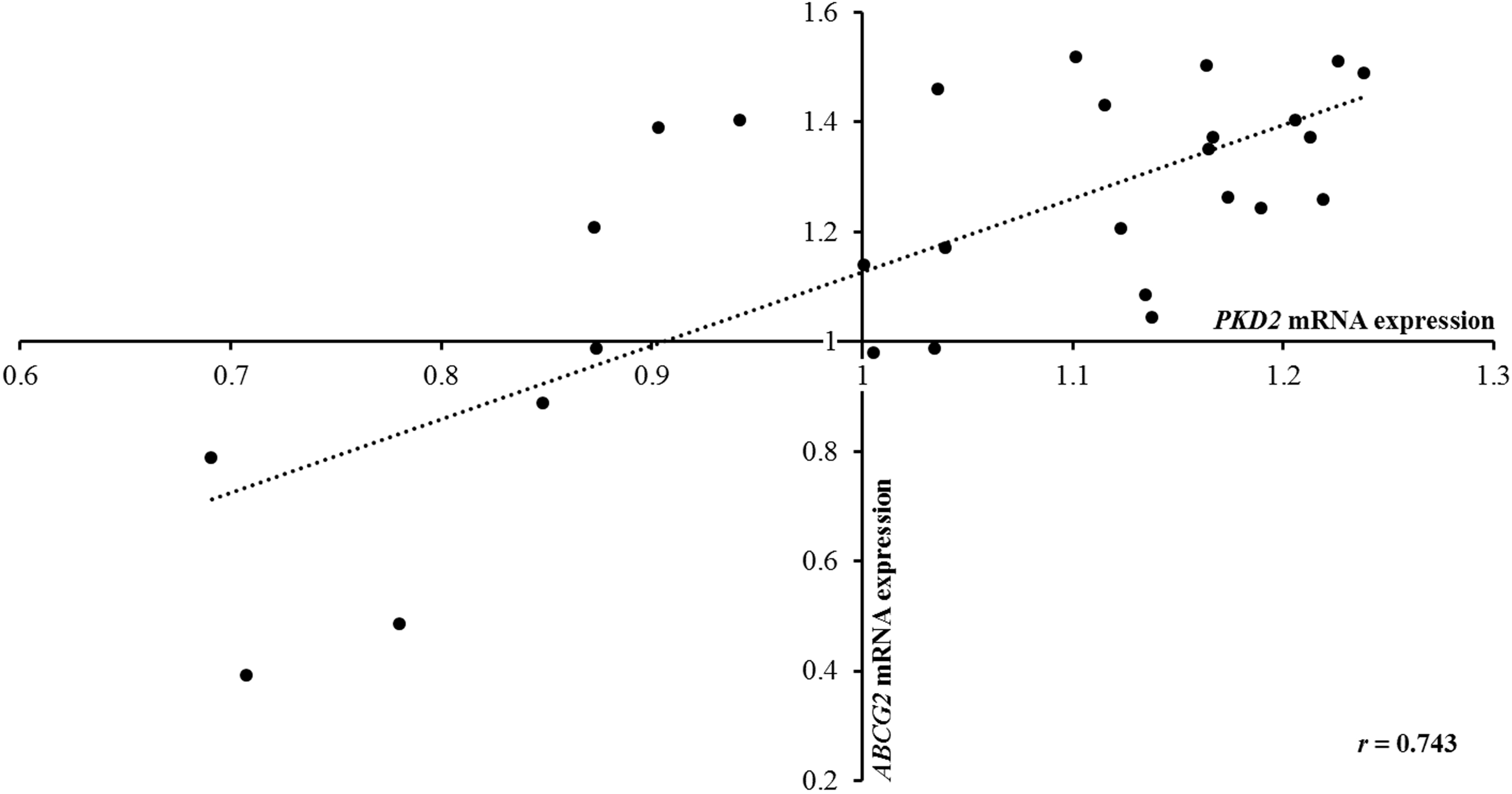
The mRNA expression correlation between *PDK2* and *ABCG2*.

To further analyze the regulation function in *PKD2* region, two SNP sets in this region were tested using enhancer enrichment analysis (Table 4). For set one, we found significant enrichments of enhancers in Huvec cell line (3.6-fold enrichment, *P* = 0.012), especially for strong enhancers (5.7-fold enrichment, *P* = 0.005) (Table 4). More significant enhancer enrichments were found for set two and those enrichments were existed in various cell lines (H1, HepG2, Huvec, HSMM, NHLF, HMEC, GM12878 and NHEK cell lines) (Table 4). All those results revealed that regulation factors in *PKD2* might played an important role in controlling the concentrations of serum urate by epistatic interacting with *ABCG2*.

**Table 4.**
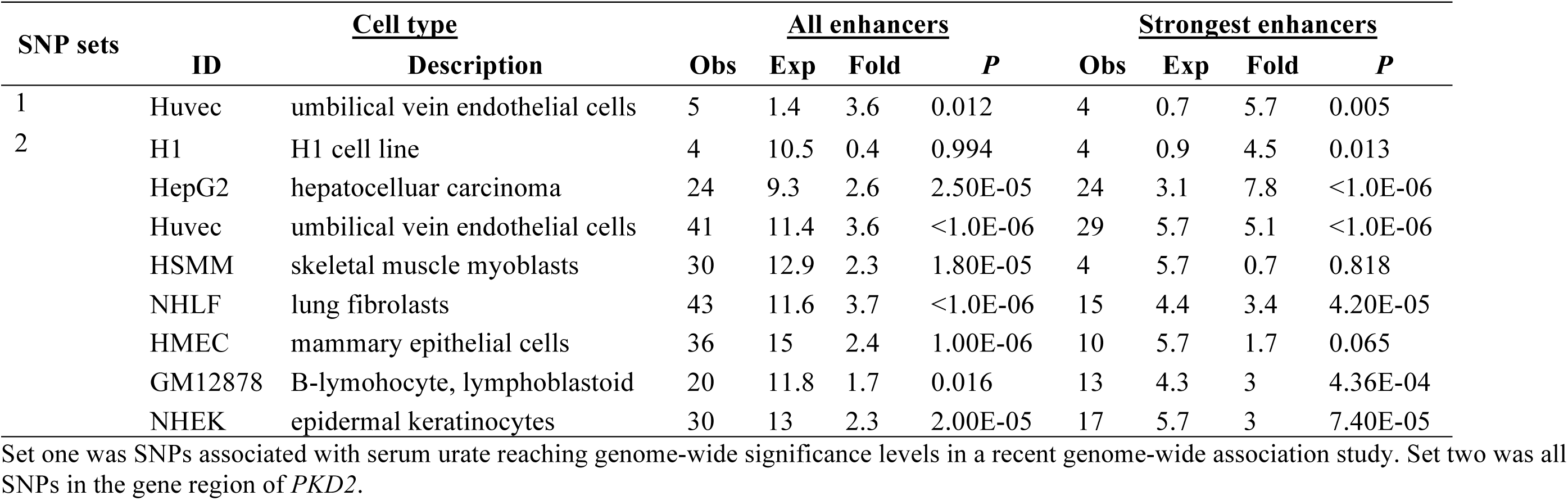
Enhancer enrichment analysis of loci in *PKD2*

| SNP sets | <u>Cell type</u> |  | <u>All enhancers</u> |  |  |  | <u>Strongest enhancers</u> |  |  |  |
| --- | --- | --- | --- | --- | --- | --- | --- | --- | --- | --- |
|  | ID | Description | Obs | Exp | Fold | *P* | Obs | Exp | Fold | *P* |
| 1 | Huvec | umbilical vein endothelial cells | 5 | 1.4 | 3.6 | 0.012 | 4 | 0.7 | 5.7 | 0.005 |
| 2 | H1 | H1 cell line | 4 | 10.5 | 0.4 | 0.994 | 4 | 0.9 | 4.5 | 0.013 |
|  | HepG2 | hepatocellular carcinoma | 24 | 9.3 | 2.6 | 2.50E-05 | 24 | 3.1 | 7.8 | <1.0E-06 |
|  | Huvec | umbilical vein endothelial cells | 41 | 11.4 | 3.6 | <1.0E-06 | 29 | 5.7 | 5.1 | <1.0E-06 |
|  | HSMM | skeletal muscle myoblasts | 30 | 12.9 | 2.3 | 1.80E-05 | 4 | 5.7 | 0.7 | 0.818 |
|  | NHLF | lung fibroblasts | 43 | 11.6 | 3.7 | <1.0E-06 | 15 | 4.4 | 3.4 | 4.20E-05 |
|  | HMEC | mammary epithelial cells | 36 | 15 | 2.4 | 1.00E-06 | 10 | 5.7 | 1.7 | 0.065 |
|  | GM12878 | B-lymphocyte, lymphoblastoid | 20 | 11.8 | 1.7 | 0.016 | 13 | 4.3 | 3 | 4.36E-04 |
|  | NHEK | epidermal keratinocytes | 30 | 13 | 2.3 | 2.00E-05 | 17 | 5.7 | 3 | 7.40E-05 |

Set one was SNPs associated with serum urate reaching genome-wide significance levels in a recent genome-wide association study. Set two was all SNPs in the gene region of *PKD2*.

### The mRNA expression correlation between *PDK2* and *ABCG2*

To further validate the potential epistatic influence in *PKD2* region, the mRNA expression correlation between *PKD2* and *ABCG2* was studied. Our result showed that *PKD2* mRNA expression was positive correlation with *ABCG2* mRNA expression (r = 0.743, *P* = 5.83e-06) (Figure 1), suggested the potential effect of regulation factor in *PKD2* on modification of *ABCG2* expression.

## Discussion

This study explored the epistatic interaction between *PKD2* and *ABCG2* affecting the pathogenesis from elevated serum urate to hyperuricemia to gout and found two SNP pairs (rs2728121:rs1481012 and rs2728121:rs2231137) significantly associated with serum urate and/or gout. The SNP pairs identified in the association study were analyzed in detail, including the analysis of association among subgroups of heterogeneity factors (such as gender, body mass index (BMI) and smoking status), functional analysis and mRNA expression analysis.

*ABCG2* gene encoded a urate transporter and contributed to serum urate concentration and gout risk. It had been proved to be one of strongest risk factor for the development of gout (Köttgen et al. 2012; Dong et al. 2015a). But the results for the association between *PKD2* and uric acid/gout is contradictory. Recent studies suggested that *PKD2* associated with uric acid and gout (Zhang et al. 2016), while other researchers had an opposite opinion (Dehghan et al. 2008). In addition, no functional experiment showed that *PKD2* directly affect the concentration of serum urate and gout risk. Therefore, this study attempted to provide more evidences to obtain a reliable conclusion.

Previous study had been suggested the important role of epistatic interaction in the 4p16.1 region regulating uric acid levels and showed a suggestive interaction pair in the gene region of *PKD2* (Wei et al. 2014). Therefore, this study focused on the epistasis effect in *PKD2* and *ABCG2* gene region. The present study showed that two epistatic interaction pairs (rs2728121:rs1481012 and rs2728121:rs2231137) were identified to associate with the concentrations of serum urate (Estimate = -14.487, *P* _int_ = 0.018 and Estimate = 9.781, *P* _int_ = 0.004, respectively) and the latter pair (rs2728121: rs2231137) was also associated with gout (*P* _int_ = 0.001) (Table 1 and 2). The results provided a new view for explaining the influence of *PKD2* on serum urate/gout. *PKD2* could contribute to the pathogenesis from elevated serum urate to hyperuricemia to gout by an indirect way as interacting with *ABCG2*. Although the direct function of *PKD2* in influencing the pathogenesis could not identified in this study, it provided a new evidence to state that *PKD2* is a urate/gout-associated gene. Our study also found that the former interaction pair could explain 0.099% of uric acid variance especially 0.493% in female, and the later interaction pair could explain 0.164% of the variance (Table 1), suggesting epistasis effect is helpful to solve the ‘missing heritability’ problem. In addition, in consistent with our previous results (Dong et al. 2015b, 2017b; Yang et al. 2016). Environment factors, such as gender and BMI modified the associations between SNP pairs and serum urate/gout in the present study (Table 3).

The mechanism of the interactions between genes were analyzed by functional analysis and mRNA expression analysis. The functional analysis showed that regulation factors enrichment into the gene region of *PKD2*, indicating the potential regulation function of *PKD2* to *ABCG2*. To further analyze this result, mRNA expression correlation between genes was used. The result suggested a positive correlation between *PDK2* mRNA expression and *ABCG2* mRNA expression, further supporting our hypothesis that *PKD2* influenced serum urate by epistatic interacting with *ABCG2*.

In conclusion, this study for the first time suggested that epistatic interactions between *PKD2* and *ABCG2* influenced serum urate concentrations and gout risk, and *PKD2* might affect the pathogenesis from elevated serum urate to hyperuricemia to gout by modifying *ABCG2*. Furthermore, our study provided a new evidence to support that *PKD2* is a urate/gout-associated gene and indicated that epistasis effect is helpful to solve the ‘missing heritability’ problem.

## Acknowledgements

General: Computational support was provided by the High-End Computing Center located at Fudan University.

Funding: This work was supported by grants from the Science and Technology Committee of Shanghai Municipality [11DJ1400100 to J.W.], International S&T Cooperation Program of China [2013DFA30870 to J.W.], Ministry of Science and Technology [2011BAI09B00 to J.W.], and Program for 2012 Outstanding Medical Academic Leader for Hejian Zou.

Author contributions: Conceived and designed the experiments: JW ZD. Performed the experiments: ZD. Analyzed the data: ZD. Contributed reagents/materials/analysis tools: JZ DZ CY YM HH HJ SJ YL. Wrote the paper: JW ZD HZ LJ.

## COMPLIANCE WITH ETHICS GUIDELINES

All authors declare that there is no conflict of interest.

All procedures followed were in accordance with the ethical standards of the responsible committee on human experimentation (institutional and national) and with the Helsinki Declaration of 1975, as revised in 2000. Informed consent was obtained from all patients for being included in the study.

## Supplementary Materials

**Figure S1. Chromatin state analysis of the *PKD2* and *ABCG2* gene region by Enlight**

**Figure S2. Chromatin state analysis of the *PKD2* and *ABCG2* gene region by UCSC genome browser**

**Table S1. Characteristics of participants in this study.** HUA, hyperuricemia. The data are shown as the mean (SD).

